# *TWIST1* controls cellular senescence and energy metabolism in mesenchymal stem cells

**DOI:** 10.1101/2020.10.11.335448

**Authors:** Chantal Voskamp, Laura A. Anderson, Wendy J. L. M. Koevoet, Sander Barnhoorn, Pier G. Mastroberardino, Gerjo J.V.M. van Osch, Roberto Narcisi

**Author notes:** **Contact corresponding author** Roberto Narcisi, Department of Orthopaedics, Erasmus MC, 3015 CN Rotterdam, the Netherlands.

## Abstract

Mesenchymal stem cells (MSC) are promising cells for regenerative medicine therapies, because they can differentiate towards multiple cell lineages. However, heterogeneity in differentiation capacity is one of the main drawbacks that limit their use clinically. Differences in the occurrence of cellular senescence and in the expression of the senescence associated secretory phenotype (SASP) in MSC populations contribute to their heterogeneity. Here, we show the involvement of *TWIST1* expression in the regulation of MSC senescence, demonstrating that silencing of *TWIST1* in MSCs increased the occurrence of senescence. These senescent MSCs had a SASP that was different from irradiation-induced senescent MSCs. In addition, metabolic evaluation performed by the Seahorse XF apparatus showed that both *TWIST1* silencing-induced and irradiation-induced senescent MSCs had a higher oxygen consumption compared to control MSCs, while *TWIST1* silencing-induced senescent MSCs had a low extracellular acidification rate compared to the irradiation-induced senescent MSCs. Overall, our data indicate how *TWIST1* regulation influences senescence in human MSCs and that *TWIST1* silencing-induced senescence is characterized by a specific expression of the SASP and the metabolic state.

## Introduction

Regenerative medicine strategies aim to regenerate tissues that have been damaged by injury or pathology. A promising cell source for regenerative medicine therapies is the multipotent progenitor cell referred to as mesenchymal stem cell (MSC). MSCs have the capacity to self-renew, and to differentiate towards multiple lineages (Pittenger et al., 1999). MSCs can be isolated from several tissues such as the bone marrow (Haynesworth et al., 1992; Pittenger et al., 1999), umbilical cord blood (Erices et al., 2000; Romanov et al., 2003), or adipose tissue (Halvorsen et al., 2000; Zuk et al., 2001). However, a limitation that hinders the clinical use of MSCs is their inter- and intra-donor variation in differentiation capacity. This heterogeneity includes the occurrence of cellular senescence (Li et al., 2017). Cellular senescence is an irreversible state in which cells undergo permanent cell cycle arrest, while the cells are still metabolically active and can secrete pro-inflammatory factors. These secreted factors are named the senescence-associated secretory phenotype (SASP) (Lunyak et al., 2017). The occurrence of the SASP is linked to the metabolic state of the cell (Dörr et al., 2013; Wiley et al., 2016). Glycolysis, which breaks down glucose into pyruvate, ATP and NADH, has been demonstrated to be increased in senescent cells (Bittles and Harper, 1984; James et al., 2015). In addition, senescent fibroblasts can have an impaired mitochondrial metabolism (Wiley et al., 2016).

Cellular senescence has been shown to reduce the differentiation capacity of umbilical cord-derived MSCs (Cheng et al., 2011) and could also be unsafe for regenerative medicine strategies, since senescent MSCs can promote tumor formation (Hochane et al., 2017; Li et al., 2015). In addition senescent cells transplanted in the knee joint of mice can induce an osteoarthritis-like phenotype showing reduced cartilage content, osteophyte formation and subchondral bone structure alterations (Xu et al., 2017). Safe and reproducible clinical use of MSCs requires a better understanding of the molecular mechanisms behind cellular senescence and their SASP profile.

Previously, we and others observed that MSC expansion was associated with the expression of the transcription factor *TWIST1* (Isenmann et al., 2009; Narcisi et al., 2015; Voskamp et al., 2020). Moreover, TWIST1 can regulate the expression of cellular senescence marker P21 in hypoxic MSC cultures (Tsai et al., 2011). To better understand the molecular mechanism behind cellular senescence in MSCs, we investigated how *TWIST1* expression regulates cellular senescence and their SASP expression. In this study we show that *TWIST1* overexpression in MSCs inhibited cellular senescence, while silencing of *TWIST1* induced cellular senescence. In addition, we show that *TWIST1* can modulate the SASP and bioenergetic profile in senescent MSCs. These results provide novel molecular insights in SASP and metabolism regulation and suggest that TWIST1 could be a target to modulate cellular senescence.

## Results and Discussion

### *TWIST1* expression is negatively associated with cellular senescence in MSCs

To determine whether *TWIST1* expression is involved in cellular senescence in human MSCs, we started by analyzing its expression in senescent MSCs. Cellular senescence was induced in MSCs by gamma irradiation (20 Gy) and confirmed by SA-β-gal staining (**Figure 1A**). *TWIST1* expression was significantly reduced in irradiated-induced senescent MSCs compared to mock irradiated MSCs (**Figure 1B**; p=0.022), indicating that *TWIST1* expression is negatively associated with cellular senescence in MSCs. Following this observation, we hypothesized that high expression of *TWIST1* is able to delay the entrance into the senescence state during passaging *in vitro*. To test this hypothesis, *TWIST1* was overexpressed in MSCs by a lentiviral-based approach. Transduction was determined by the percentage of GFP positive cells (>65% transduced cells; **Fig EV1**) and overexpression confirmed by qPCR analysis (103-fold increase compared to empty vector control; **Figure 1C**). Control and *TWIST1* overexpressing MSCs were then serially passaged for 11 days, followed by SA-β-gal analysis (**Fig EV2**). *TWIST1* overexpressing MSCs showed 15% SA-β-gal low positive cells and 0.4% SA-β-gal high positive cells, while empty vector control cells had 52% SA-β-gal low positive cells (p<0.001) and 2% high positive cells (p=0.052; **Figure 1D**). These results suggest that *TWIST1* expression can inhibit cellular senescence in MSC.

**Figure 1.**
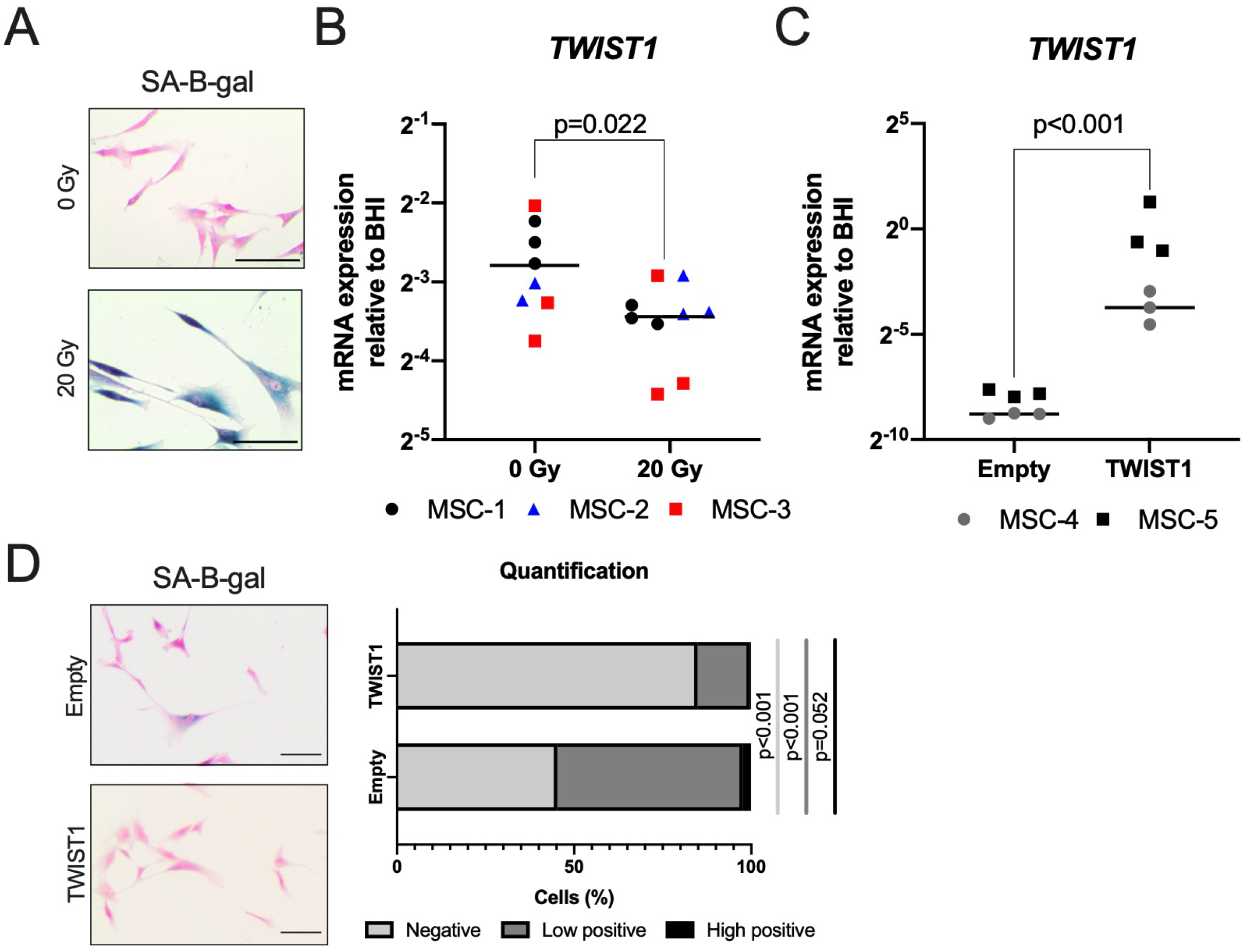
*TWIST1* expression is negatively associated with senescence-associated β-galactosidase. (A) Representative images of senescence-associated β-galactosidase (SA-β-gal) staining counter stained with pararosaniline of MSCs 7 days after gamma irradiation with 0 or 20 Gy. Scale bar represent 100 µm. N=9, 3 donors with 3 replicates per donor. (B) *TWIST1* mRNA levels of MSCs 7 days after gamma irradiation with 0 or 20 Gy. Data show individual data points and grand mean with N=8 (0 Gy) or N=9 (20 Gy), 3 donors with 2-3 replicates per donor, linear mixed model. (C) *TWIST1* mRNA levels of MSCs transduced with an empty overexpression lentiviral construct (Control) or a TWIST1 overexpression lentiviral construct (TWIST1) after 11 days of expansion. Data show individual data points and grand mean with N=6, 2 donors with 3 replicates per donor, linear mixed model. (D) Left panel, representative images of SA-β-gal staining counter stained with pararosaniline of MSCs transduced with an empty overexpression lentiviral control construct (Empty) or a TWIST1 overexpression lentiviral construct (TWIST1) after 11 days of expansion. Right panel, quantification of SA-β-gal staining. Bars show grand mean of percentage of SA-β-gal negative, low positive and high positive cells. N=4, 2 donors with 2 replicates per donor, linear mixed model.

It has been reported that *TWIST1* expression suppresses senescence in lung and breast cancer cells (Burns et al., 2013; Nayak et al., 2017; Tran et al., 2012). In MSCs, a high *TWIST1* expression has been associated with rapid cell growth and a high proliferation capacity of MSCs (Boregowda et al., 2016; Isenmann et al., 2009; Voskamp et al., 2020). Our data indicate a direct link between *TWIST1* expression and cellular senescence in MSCs.

### *TWIST1* silencing induces cellular senescence with a specific senescence associated secretory phenotype in MSCs

To elucidate whether cellular senescence can be induced via *TWIST1* modulation, *TWIST1* expression was silenced in MSCs using a siRNA approach. After 24 h of *TWIST1* siRNA treatment (siTWIST1-MSCs) *TWIST1* mRNA levels were 53% reduced (p=0.035) compared to scramble controls (**Fig EV3A**) and after 4 passages siTWIST1-MSCs showed 64% knockdown of *TWIST1* mRNA levels (p<0.001; **Fig 2A**). After 4 passages, *TWIST1* silencing increased the expression of cell cycle inhibitors and senescence markers *P16* (6.5-fold, p<0.001) and *P21* (2.1-fold, p=0.060; **Fig 2B**). The expression of *P16* was already induced after 24 h of *TWIST1* silencing (1.8-fold; p=0.015; **Fig EV3B**). No differences in *P21* expression were observed after 24 h of *TWIST1* silencing (**Fig EV3C**). In addition, after 4 passages, *TWIST1* silencing increased SA-β-gal activity in MSCs (**Fig 2C and Fig EV2E**) and decreased cell expansion (**Fig 2D**), overall indicating that *TWIST1* knockdown induces senescence growth arrest. Since the SASP can drive chronic inflammation and thereby contribute to age-related diseases such as osteoarthritis and cancer (reviewed in: Loeser et al., 2016; Zhu et al., 2014), we determined the expression of the SASP-related genes *IL6, IL10, IL1B, MMP3, IL8, CCL2* and *VEGFA* in siTWiST1-MSCs. siTWIST1-MSCs expressed higher levels of *CCL2* and *IL1B* compared to control condition (3.3-fold p=0.008, 7.4-fold p=0.008, respectively; **Fig 2E**). Interestingly, the expression of *IL6, IL10, MMP3* and *VEGFA* were not significantly affected by *TWIST1* silencing, and *IL8* was even significantly decreased (p=0.291, p=0.077, p=0.087, p=0.912, p<0.001, respectively; **Fig 2E**). These results indicate that senescence was induced in MSCs by *TWIST1* knockdown, but generating a ‘non-classical’ SASP profile.

**Figure 2.**
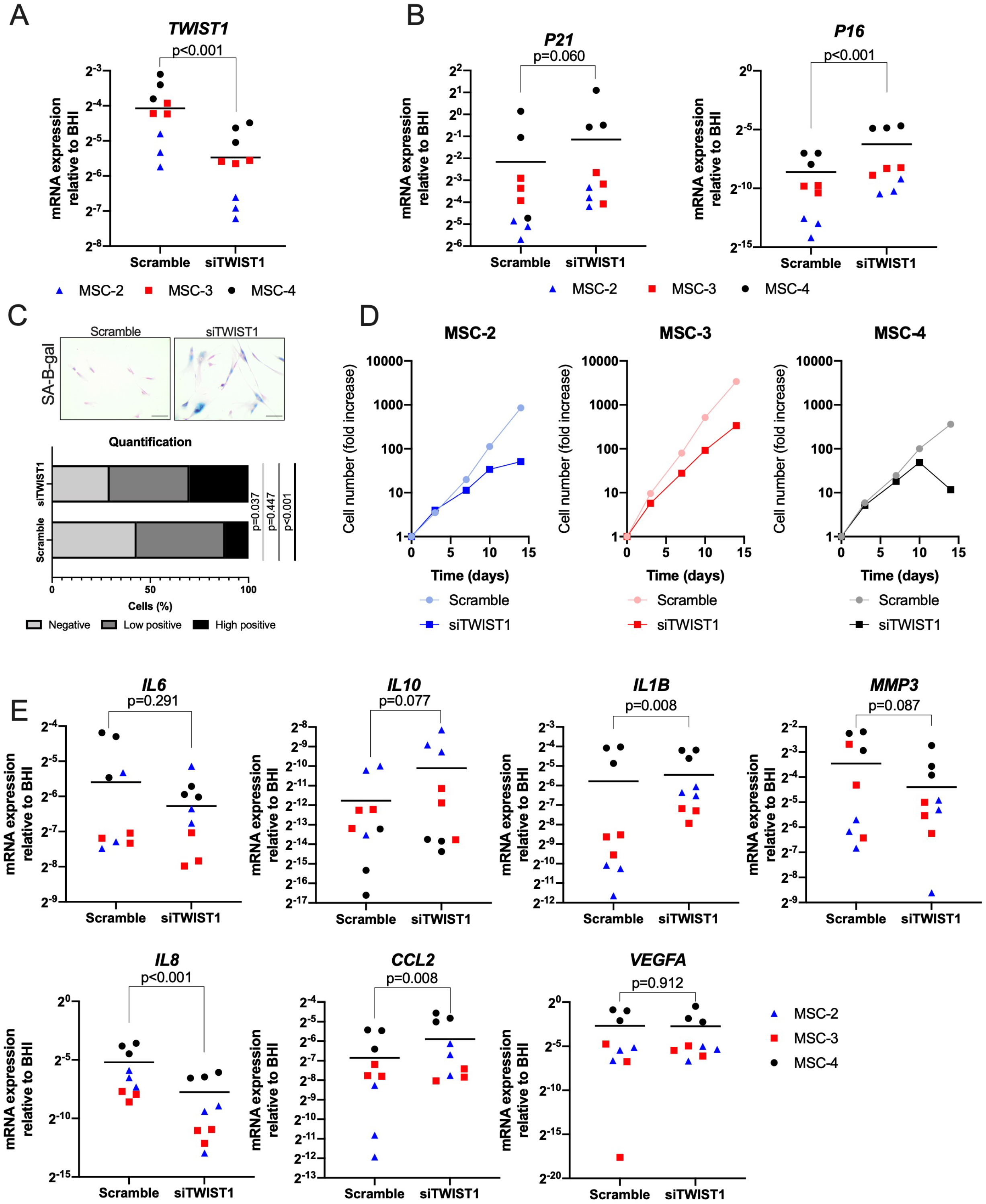
*TWIST1* silencing induces cellular senescence in MSCs with a specific SASP mRNA expression profile. (A) *TWIST1* mRNA levels in MSCs treated for 4 passages with scramble siRNA (Scramble) or siRNA against TWIST1 (siTWIST1). N=9, 3 donors with 3 replicates per donor, linear mixed model. (B) *P16* and *P21* mRNA levels in MSCs treated for 4 passages with scramble siRNA (Scramble) or siRNA against TWIST1 (siTWIST1). N=9, 3 donors with 3 replicates per donor, linear mixed model. (C) Quantification of senescence associated beta galactosidase (SA-β-gal) staining based on intensity (negative, low positive and high positive) of MSCs treated for 4 passages with scramble siRNA (Scramble) or siRNA against TWIST1 (siTWIST1). N=6, 3 donors with 2 replicates per donor, linear mixed model. (D) Cell number data during expansion of MSCs treated with scramble siRNA (Scramble) or siRNA against TWIST1 (siTWIST1) at day 0, 3, 7, 10 and 14 of treatment, N=3 donors. (E) *IL6, IL10, IL1B, MMP3, IL8, CCL2* and *VEGFA* mRNA levels in MSCs treated for 4 passages with scramble siRNA (Scramble) or siRNA against TWIST1 (siTWIST1). N=9, 3 donors with 3 replicates per donor, linear mixed model. Graphs show individual data points and grand mean.

The mechanism by which the SASP related genes are regulated in senescent cells is not fully understood. It is, however, known that mitochondria can induce SASP expression via increased reactive oxygen species (ROS) and JNK activation (Vizioli et al., 2020). Interestingly, cellular senescence with a different SASP profile has previously been reported in mitochondrial dysfunctional senescence (MiDAS) (Wiley et al., 2016). Cells with MiDAS appeared to have a SASP expression profile similar to siTWIST1-MSCs: they did not express *IL-6* and *IL8* and had an increased expression of *IL-10* (Wiley et al., 2016). In addition, *TWIST1* downregulation was demonstrated to promote mitochondrial dysfunction in lung cancer cells (Seo et al., 2014) and adipocytes (Lu et al., 2018). Overall, these data suggest that *TWIST1* silencing might induce cellular senescence in MSCs via mitochondrial dysfunction.

### *TWIST1* silencing alters MSC bioenergetics

To study if *TWIST1* silencing-induced senescence is induced via mitochondrial dysfunction, we determined the bioenergetic profile in siTWIST1-MSCs using a Seahorse XF-24 Extracellular Flux Analyzer. We measured the oxygen consumption rate (OCR) reflecting cellular respiration followed by subsequent measurement after injection of mitochondrial toxins: oligomycin, FCCP and antimycin A (see materials and methods and **Fig EV4A**). We first identified the optimal cell density (30,000 cells/well; **Fig EV4B**) and the ideal concentration of FCCP (2.0 μM**; Fig EV4C**) to detect OCR in human MSCs. Then, we observed a significant increase in basal respiration levels in siTWIST1-MSCs compared to scramble controls (p=0.011; **Fig 3A-C**). In addition, siTWIST1-MSCs showed a higher maximum OCR, proton leak, Adenosine Tri-Phosphate (ATP) production and spare respiratory capacity compared to scramble control cells (p=0.001, p=0.006, p=0.002, p=0.002; **Fig 3D-G**). No differences in non-mitochondrial respiration were observed between scramble control cells and siTWIST1-MSCs (p=0.251; **Fig 3H**). These data indicate that that *TWIST1* silencing induces changes in the mitochondrial function in MSCs. To determine if an increased mitochondrial respiration is specific for *TWIST1* silencing-induced senescent MSCs or whether it is common for senescent MSCs, we determined the OCR in irradiated induced senescent MSCs. Similar to *TWIST1* silencing-induced senescent MSCs, irradiation-induced senescent MSCs showed higher basal OCR, maximum OCR, proton leak, ATP production and spare respiratory capacity compared to non-irradiated control cells (p<0.001, p=0.050, p=0.019, p<0.001, p=0.032; **Fig EV5A-F**), and no differences in non-mitochondrial respiration (p=0.256; **Fig EV5G**). These data suggest that both *TWIST1* silencing-induced and irradiation-induced senescent MSCs have an increased OCR. Previously, cellular senescence has been associated with an increased OCR in fetal lung cells (Quijano et al., 2012). The increased OCR in senescent MSCs can be due to an increase in mitochondrial respiration or to an increase in mitochondrial mass. An increase in mitochondrial mass in senescent cells has been reported before in fibroblasts (Correia-Melo et al., 2016; Lee et al., 2002) and can increase the levels of mitochondrial-derived reactive oxygen species (ROS) which can cause an increase in proton leak (Brookes, 2005). In both *TWIST1* silencing-induced and irradiation-induced senescent MSCs we indeed observed an increased proton leak (**Fig 3** and **Fig EV5**), suggesting that the senescent MSCs have dysfunctional mitochondria. Dysfunctional mitochondria can trigger cellular senescence (Wiley et al., 2016), and removal of mitochondria in senescent cells has been shown to reduce the senescence phenotype (Correia-Melo et al., 2016), indicating that mitochondria can induce cellular senescence, but also play a key role in the maintenance of the senescence phenotype. Despite the difference in the SASP phenotype, both *TWIST1* silencing-induced and irradiation-induced senescent MSCs showed an increased mitochondrial respiration.

**Figure 3.**
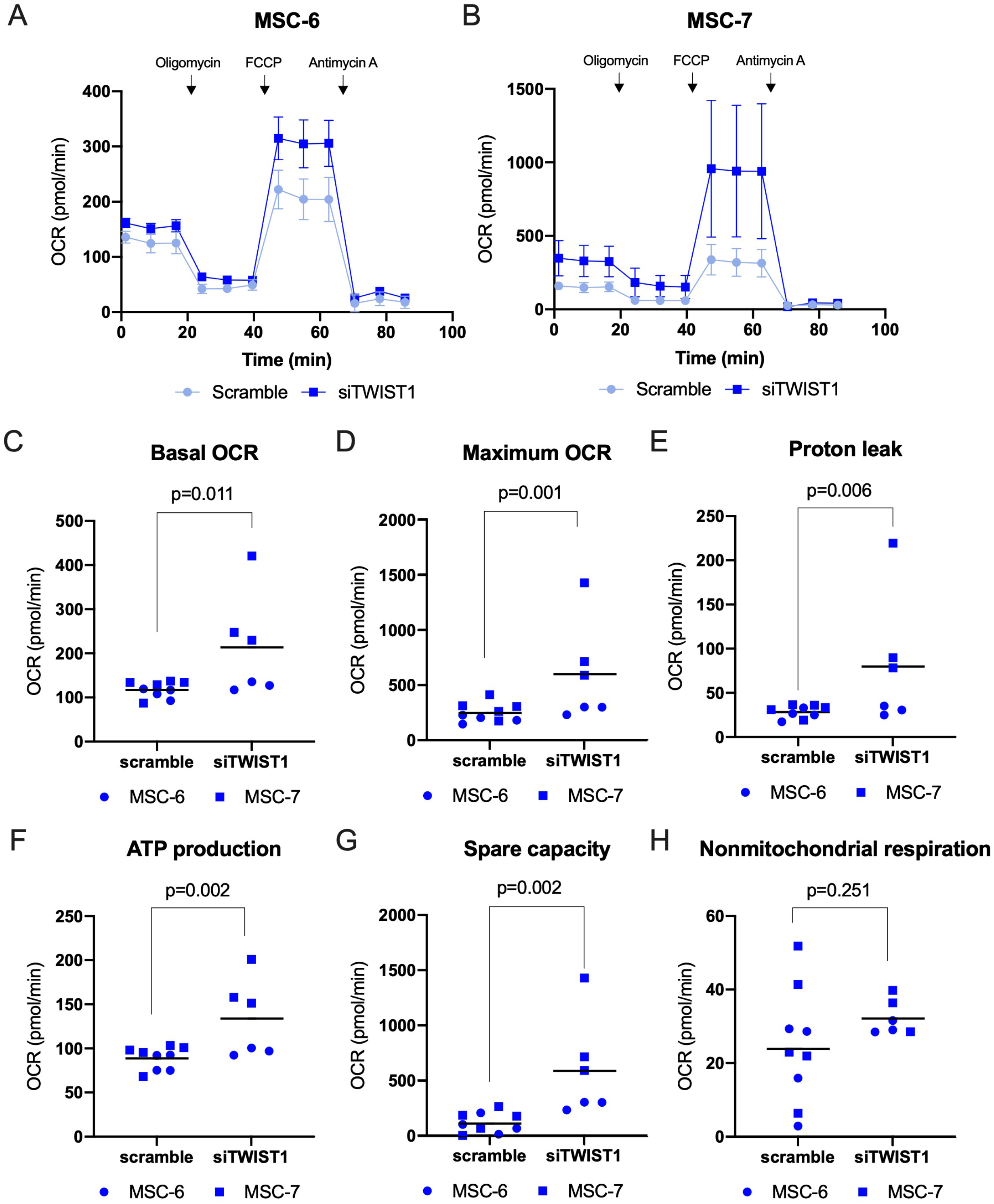
Increased oxygen consumption rate (OCR) in *TWIST1* silenced MSCs. (A-B) Graph shows the OCR in MSCs treated with a scramble or TWIST1 siRNA at basal level and after addition of oligomycin, FCCP and antimycin A in two different donors MSC-6 (A) and MSC-7 (B). Values represent mean with SD, N=3-5 replicates per donor. (C-H) Graphs show calculated values for basal OCR (C), maximum OCR (D), proton leak (E), ATP production (F), spare capacity (G) and non-mitochondrial respiration (H) in MSCs treated with scramble or TWIST1 siRNA. N=6-9, 2 donors with 3-5 replicates per donor, linear mixed model. Graphs show individual data points and grand mean.

In addition to mitochondrial respiration, glycolysis plays an important role in MSC energy metabolism (Pattappa et al., 2011). Cellular senescence has been associated with an increased glycolytic capacity after *in vitro* expansion (Bittles and Harper, 1984). As a measure of glycolytic flux in siTWIST1-MSCs, we analyzed the extracellular acidification rate (ECAR). No significant differences in ECAR were observed between scramble control cells and siTWIST1-MSCs (**Fig 4A-C**), indicating that *TWIST1* silencing does not alter the glycolytic flux in MSCs. Irradiated MSCs had higher ECAR compared to control MSCs (**Fig EV6**), confirming earlier published data in fibroblasts (James et al., 2015). These data suggest that the glycolytic capacity is unaltered in siTWIST1-MSCs, in contrast to irradiation induced senescent MSCs. Overall, these data show that depending on the inducer of cellular senescence in MSCs, senescent MSCs can have a different bioenergetic profile.

**Figure 4.**
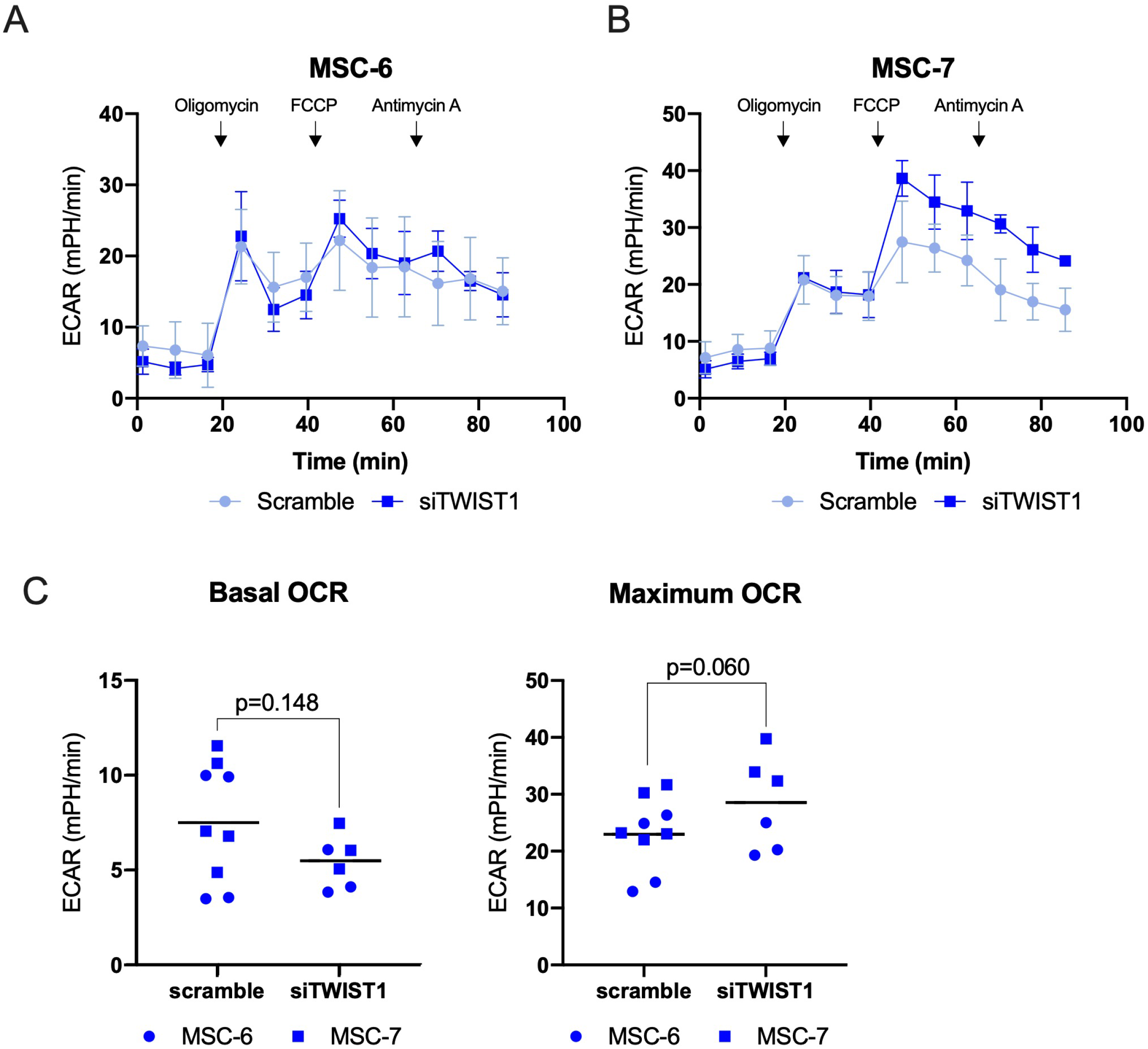
*TWIST1* silencing did not increases extracellular acidification rate (ECAR) in MSCs. (A-B) Graph shows the ECAR in MSCs treated with a scramble or TWIST1 siRNA at basal level and after addition of oligomycin, FCCP and antimycin A in two different donors MSC-6 (A) and MSC-7 (B). Values represent mean with SD, N=3-5 replicates per donor. (C) Graphs show ECAR values for basal OCR and maximum OCR in MSCs treated with scramble or TWIST1 siRNA. N=6-9, 2 donors with 3-5 replicates per donor, linear mixed model. Graphs show individual data points and grand mean.

In summary our study provides novel insights in the function of *TWIST1* in the regulation of cellular senescence in MSCs. Furthermore, the phenotype of these senescent cells differs from irradiation-induced senescent cells in the expression of the SASP expression and the bioenergetics (**Fig. 5**), highlighting that senescent MSCs can be heterogeneous. Besides, our results suggest that reduction of *TWIST1* expression might drive aging phenotypes of MSCs.

**Figure 5.**
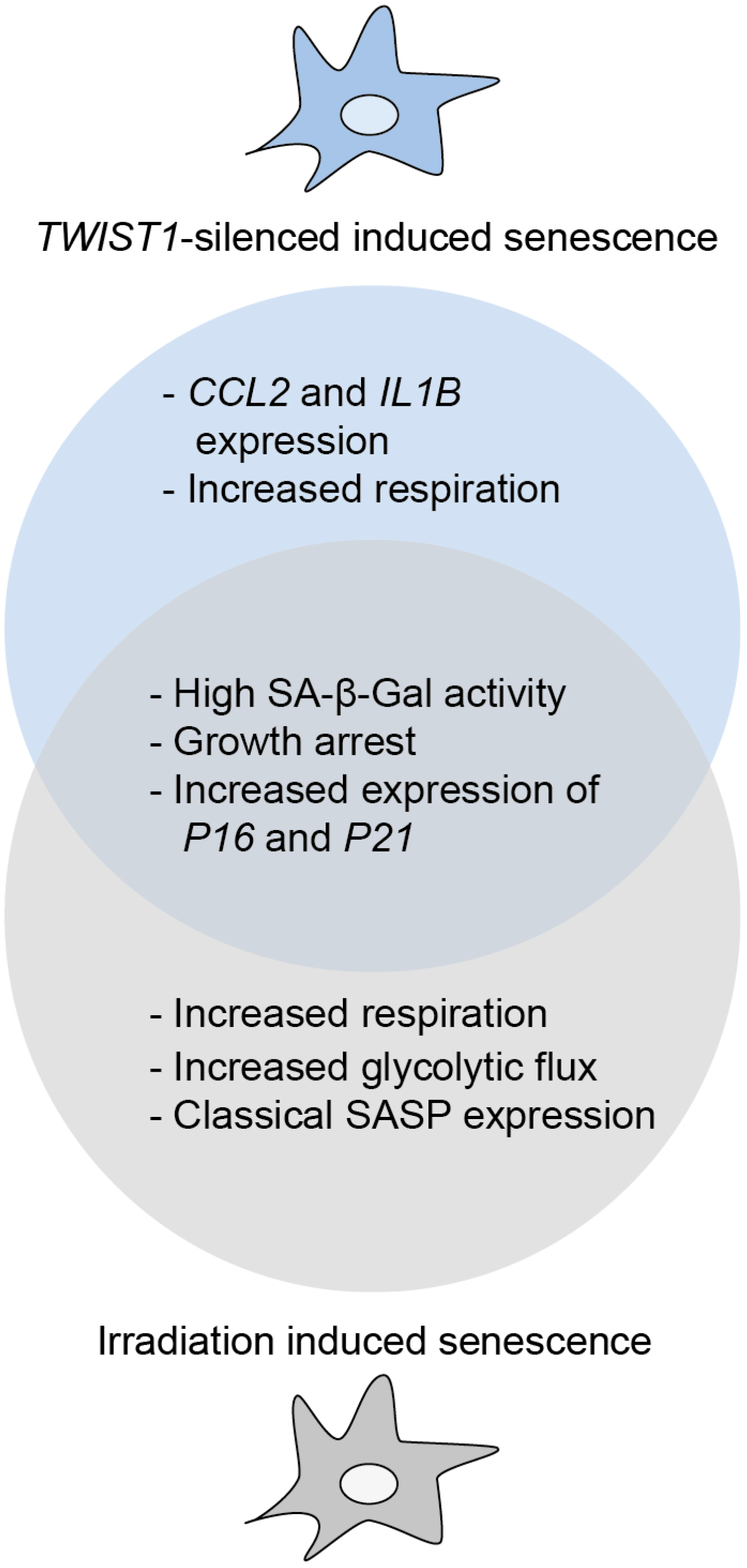
Schematic overview of the characteristics of *TWIST1* silencing-induced senescence and irradiation-induced senescence. Both *TWIST1* silencing-induced and irradiation-induced senescence stimulated senescence-associated beta-galactosidase (SA-β-gal) activity in MSCs, decreased their expansion rate and induced high levels of *P16* and *P21* mRNA. *TWIST1* silencing-induced senescence, however, increased oxygen respiration in MSCs, increased the expression of the SASP related genes *CCL2* and *IL1B*, and lack the expression of *IL8, IL6, IL10, MMP3* and *VEGFA*. Irradiation-induced senescent MSCs showed increased oxygen respiration, increased glycolytic flux and classical SASP expression.

## Materials and Methods

### Cell culture

Mesenchymal stem cells (MSC), were isolated as previously described (Knuth et al., 2018) from leftover iliac crest bone chip material obtained from patients undergoing cleft palate reconstructive surgery (MEC-2014-16; 9-13 years old). MSCs were expanded in αMEM medium (Gibco, Paisley, UK) containing 10% fetal calf serum (Gibco, selected batch 41Q2047K), 1.5 μg/mL fungizone (Invitrogen, California), and 50 µg/mL gentamicin (Gibco, Carlsbad, California), 0.1 mM Ascorbic acid (Sigma-Aldrich) and 1 ng/mL FGF2 (Instruchemie, Delftzijl, The Netherlands). MSCs were cultured at a density of 2,300 cells/cm^2^ at 37°C and 5% CO_2_. Cells were trypsinized and refreshed twice a week. Depending on the assay and the experimental plan, passage-3 (P3) to passage-7 (P7) cells have been used.

### *TWIST1* silencing

To study whether silencing of *TWIST1* induced cellular senescence, MSCs in a low passage (P3-P4) were used for this experiment. MSCs were seeded at a density of 2,300 cells/cm^2^ and cultured for 24 h in standard expansion medium. Next, the cells were treated with 15 nM TWIST1 (4390824, Ambion) or scramble (4390843, Ambion) siRNA in combination with Lipofectamine RNAMAX Transfection Reagent (1:1150; Invitrogen) and optiMEM (1:6; Gibco) or were left untreated. The treatment was repeated every 3-4 days for 13-14 days.

### Lentiviral constructs and virus generation

To study the effect of *TWIST1* overexpression on MSC senescence, lentiviral constructs of tetracycline-inducible expression of TWIST1 and green fluorescent protein (GFP) were used. TWIST1 cDNA was cloned into a lentiviral construct under the control of the tetracycline operator. The GFP lentiviral vector was a gift from Marius Wernig’s laboratory (Stanford School of Medicine, CA; Addgene plasmid # 30130). An empty lentiviral construct was used as control. Third generation lentiviral particles with a VSV-G coat were generated in HEK293T cells. HEK293T cells were cultured in DMEM HG glutamax (Life Technologies, Paisley, UK) containing 10% fetal calf serum, 1 mM sodium pyruvate (Life Technologies, Paisley, UK) and non-essential amino acids (1:100; Life Technologies, Grand Island, USA) and seeded at Poly-L-Ornithine coated plates at a density of 5 x 10^6^ cells per 10 cmØ dish. After 24 h cells were transfected with lentiviral packaging vectors PMDL (5 μg per 10 cmØ dish), RSV (2.5 μg per 10cmØ dish) VSV- (2.5 μg per 10 cmØ dish) and one of the experimental inserts; rtTA, TWIST1, GFP or an empty vector (10 μg per 10 cmØ dish) using polyethylenimine (1:166). Medium was refreshed 6 h post-transfection. Viral supernatants were filtered through a 0.45 µm filter 24 h following the last medium refreshment and stored at −80°C until use.

### Lentiviral transduction

To study whether *TWIST1* overexpression inhibited cellular senescence, MSCs in a high passage (P7) were used for this experiment. The transduction efficiency was determined in MSCs by titration of the GFP lentivirus construct using different virus concentrations, 1:1:1, 1:1:3 and 1:1:8 (GFP: rtTA: MSC expansion medium). After transduction of the cells for 16 h, cells were washed with PBS and fresh expansion medium was added with 2 μg/ml doxycycline (Sigma Aldrich). The transduction efficiency was assessed by analysis of the percentage of GFP positive cells using fluorescent microscopy and flow cytometry. For flow cytometry analysis, GFP transduced MSCs were fixed in 2% formaldehyde (Fluka) and filtered through 70-µM filters. Untransduced MSCs were used as a negative control. Samples were analyzed by flow cytometry using a BD Fortessa machine (BD Biosciences). The data were analyzed using FlowJo V10 software. Both fluorescent microscopy and flow cytometry showed that 65% of the cells were positive for GFP using a concentration of 1:1:1 lentivirus (**Fig EV1**), indicating that the MSCs were effectively transduced.

### mRNA analysis

For each experiment involving RNA evaluation, the medium was renewed 24 h before harvesting the cells. MSCs were washed with PBS and lysed in RLT with 1% β-mercaptoethanol, subsequently RNA was isolated from the cells using the RNeasy micro kit (Qiagen, Hilden, Germany) according to manufactures’ instructions. cDNA was synthesized using the RevertAid First-Strand cDNA Synthesis Kit (Thermo Fisher Scientific, Vilnius, Lithuania). Real-time polymerase chain reactions were performed with TaqMan Universal PCR MasterMix (Applied Biosystems) or SYBR Green MasterMix (Fermentas) using a CFX96TM PCR detection system (Bio-Rad). Primers are listed in **Table EV1** and housekeeping genes *GAPDH, HPRT1* and *RPS27A* were chosen for their stability in MSCs and the best housekeeping index (BHI) was calculated according to the (Ct^GAPDH^ * Ct^HPRT^ * Ct^RPS27A^)^1/3^ formula. The relative gene expression was calculated with the ΔΔCt method.

### Senescence-associated beta-galactosidase staining

Cells were washed twice with PBS and fixed with 0.5% glutaraldehyde and 1% formalin in Milli-Q water. Then the cells were washed with Milli-Q water and incubated for 24 h at 37°C with freshly made X-gal solution (0.5% X-gal, 5 mM potassium ferricyanide, 5 mM potassium ferrocyanide, 2mM MgCl2, 150mM NaCl, 7mM C6H8O7, 25mM Na2HPO4). Cells were counterstained with pararosaniline (1:25 in Milli-Q water) and detected with bright field microscopy. For each condition two independent researchers blinded to the experimental plan, scored at least 300 cells as ‘negative’, ‘low positive’, or ‘high positive’ (**Fig EV2**).

### Bioenergetics Assays

Mitochondrial respiration was measured as oxygen consumption rate (OCR) using a XF-24 Extracellular Flux Analyzer (Seahorse Bioscience) as previously described (Milanese et al., 2019). MSCs were seeded at a density of 3 x 10^4^ cells/well on Seahorse plates. Optimal cell densities were determined experimentally to ensure a proportional response to FCCP (**Fig EV4B-C**). 24 h after cell seeding, the medium was changed to unbuffered DMEM (XF Assay Medium-Agilent Technologies, Santa Clare, Ca, USA) with 2 mM glutamine, 10 mM glucose and 1 mM sodium pyruvate and incubated 1 h at 37° C in the absence of CO_2_. Three baseline measurements were performed, followed by subsequent measurements after injections of mitochondrial toxins 1.0 µM oligomycin (ATP-synthase inhibitor), 2.0 µM fluoro-carbonyl cyanide phenylhydrazone (FCCP, oxidative phosphorylation uncoupler) and 1 µM antimycin A (complex III inhibitor). Medium and reagents were adjusted to pH 7.4 according to manufacturer’s procedure. Non-mitochondrial respiration, basal respiration, proton leak, ATP production, maximal respiration and spare capacity were calculated as indicated in **Fig EV4A**: The non-mitochondrial respiration was defined as the average OCR values after antimycin A injection; basal respiration was calculated as difference between basal respiration and respiration measured after antimycin A; proton leak was calculated as difference between respiration measured after oligomycin and respiration measured after antimycin A; ATP production was calculated as difference between baseline respiration and respiration measured after oligomycin injection; maximal respiration was calculated as difference between respiration after FCCP and respiration measured after antimycin A; spare capacity was defined as difference between respiration after FCCP and baseline respiration.

## Data analysis

Results are statistically analyzed using PSAW statistics 20 software (SPSS Inc., Chicago, IL, USA). The normal distribution of the data was determined using the Kolmogorov-Smirnov test. When necessary the data was a log transformed to meet the normal distribution criteria. An unpaired t-test or a linear mixed model was applied, in this model the conditions were considered as fixed parameters and the donors as random factor. P-values less than 0.05 are considered as statistically significant. The grand mean is determined by calculating the mean of the donor means with 2-6 replicates per donor.

## Acknowledgements

The authors would like to thank Andrea Lolli for advice on the *TWIST1* silencing protocol, Eric Farrell and Janneke Witte-Bouma for access to their source of MSCs, Nicole Kops and Arielle Molina Rakos for technical assistance with the senescence-associated beta-galactosidase staining and quantification, the Marius Wernig’s laboratory (Stanford School of Medicine, CA) for providing the GFP overexpression construct and the lentiviral packaging constructs and the FACS sorting facility at the Erasmus MC for support with the BD Fortessa machine. This research was financially supported by the Dutch Arthritis Society (ReumaNederland; 16-1-201) and by a TTW Perspectief grant from NWO (William Hunter Revisited; P15-23). This study is part of the Medical Delta RegMed4D program and the Erasmus Postgraduate School Molecular Medicine.

## Author Contribution

CV: conception and design, collection of data, data analysis and interpretation, manuscript writing and final approval of manuscript. LA: design, collection of data, data analysis and interpretation of the cellular senescence analysis experiments, manuscript editing and final approval of manuscript. WK: collection of data, data analysis, manuscript editing and final approval of manuscript. SB: design, collection of data, data analysis and interpretation of the metabolic experiments and manuscript editing and final approval of manuscript. PM: conception and design and data interpretation of the metabolic experiments, manuscript editing and final approval of manuscript. GO and RN: conception and design, data analysis and interpretation, manuscript editing and final approval of manuscript.

## Data availability

All dataset generated for this study are available on request to the corresponding author.

## Expanded view Figures

**Fig EV1. Related to Fig1.**
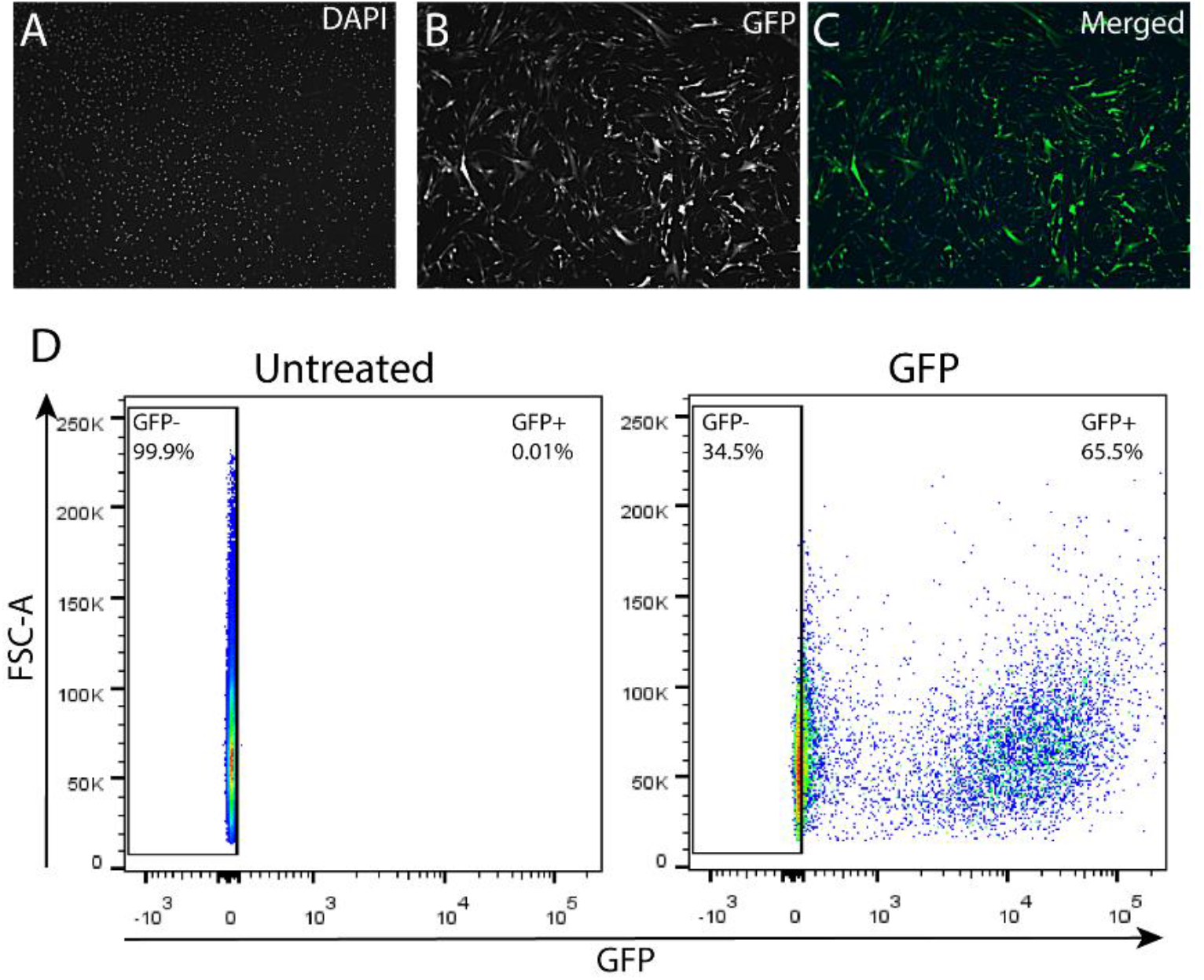
More than 65% of the MSCs were transduced using a GFP overexpression lentivirus. (A-C) Representative fluorescent microscope images of GFP lentivirus transduced MSCs stained with DAPI (nuclei), N=1 donor. (D) Flowcytometry graphs of untreated MSCs and MSCs transduced with the GFP lentivirus, N=1 donor.

**Fig EV2. Related to Fig1 and Fig 2.**
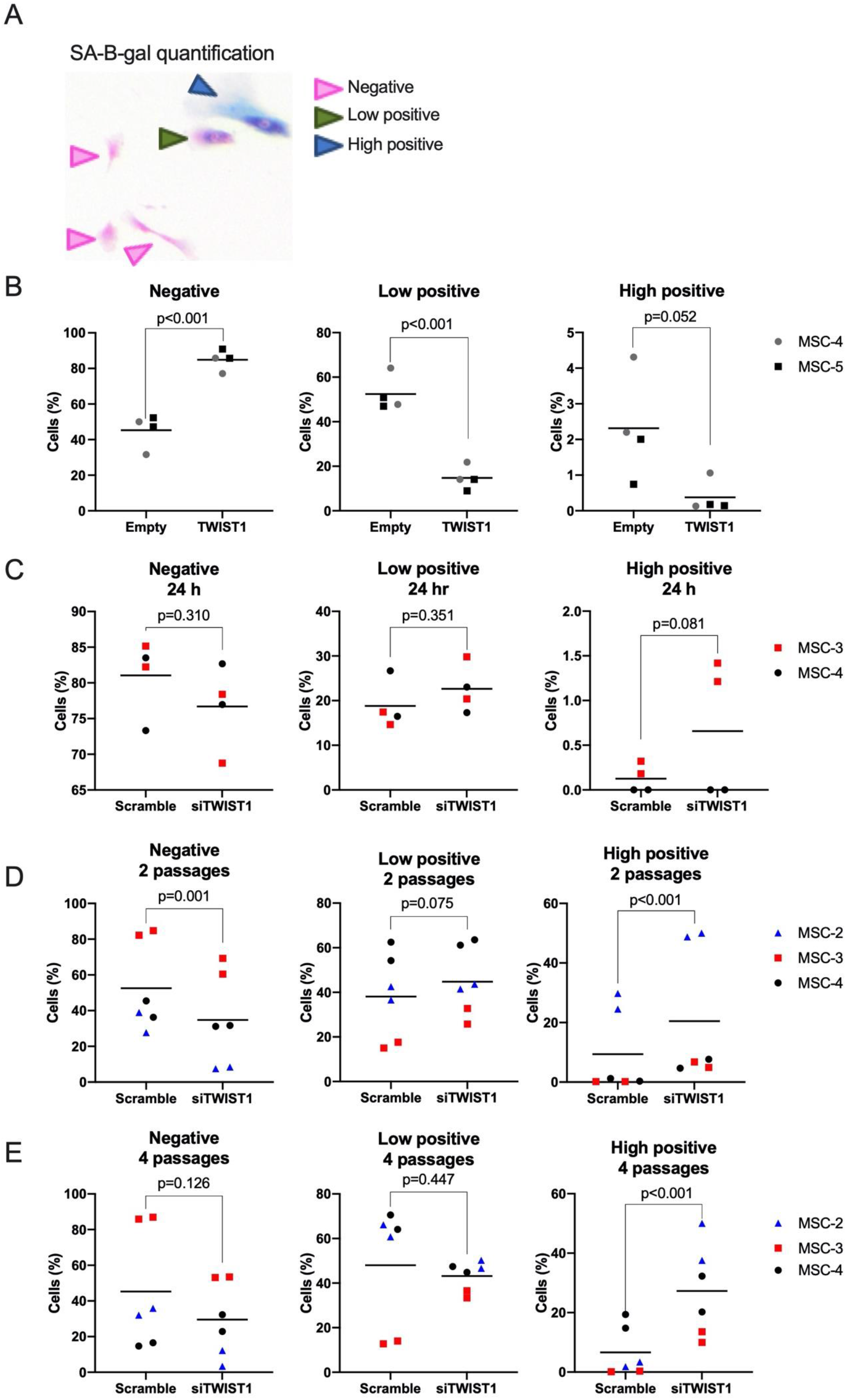
Quantification of senescence-associated β-galactosidase staining in low positive, high positive or negative cells. (A) MSCs were categorized as negative for senescence-associated β-galactosidase (SA-β-gal) staining if no blue staining was detected in the cells (pink arrow). MSCs were categorized low positive for SA-β-gal staining if cells show partial cytoplasmic staining (green arrow). MSCs were categorized as high positive for SA-β-gal staining if cells showed complete cytoplasmic staining (blue arrow). (B) SA-β-gal quantification of MSCs transduced with an empty overexpression lentiviral construct (Empty) or a TWIST1 overexpression lentiviral construct (TWIST1) after 11 days of expansion. N=4, 2 donors with 2 replicates per donor. (C-E) SA-β-gal quantification of MSCs treated for 24 h (C), 2 passages (D) or 4 passages (E) with scramble siRNA (Scramble) or siRNA against TWIST1 (siTWIST1). N=4-6, 2-3 donors with 2 replicates per donor. (B-E) Graphs show individual data points and grand mean of percentage of SA-β-gal negative (left), low positive (middle panel) and high positive (right panel) cells.

**Fig EV3. Related to Fig 2.**
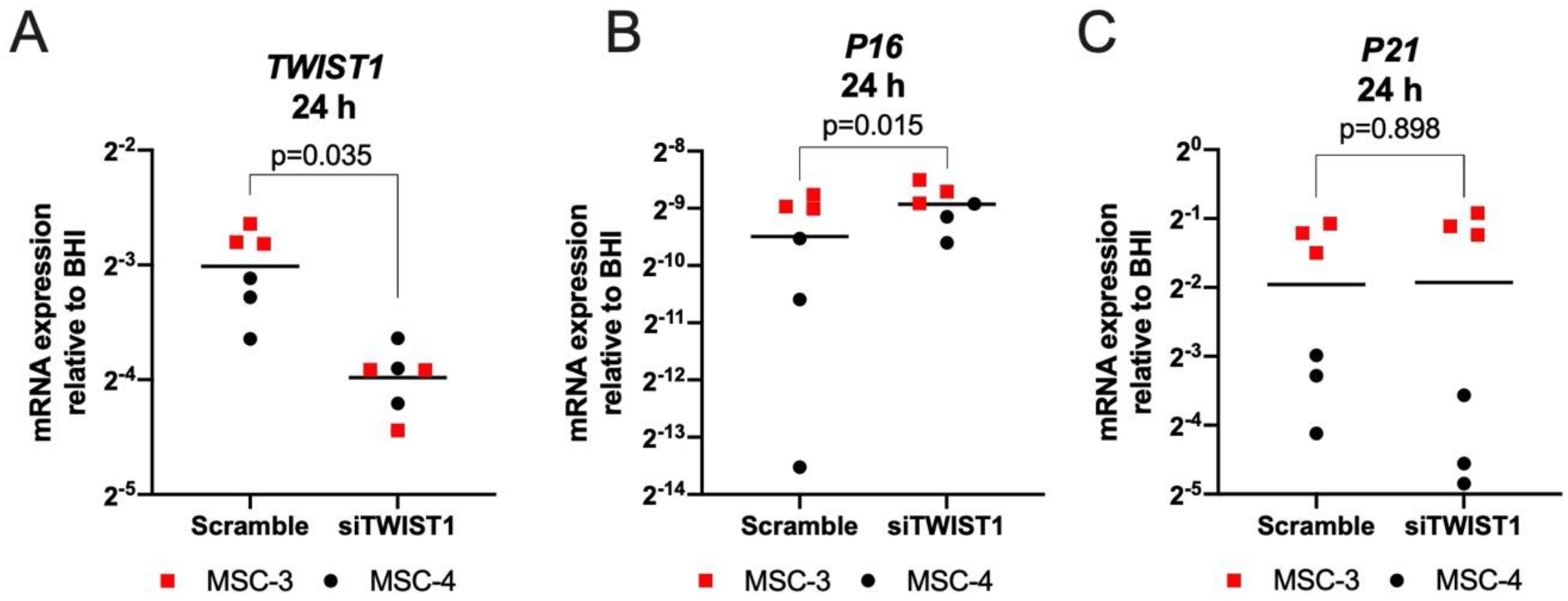
Senescence markers after 24 h and 2 passages of *TWIST1* silencing in MSCs. (A-C) *TWIST1* (A), *P16* (B), and *P21* (C) mRNA levels in MSCs treated for 24 h with scramble siRNA (Scramble) or siRNA against TWIST1 (siTWIST1). N=6-9, 2-3 donors with 3 replicates per donor, linear mixed model. Graphs show individual data points and grand mean.

**Fig EV4.Related to Fig3.**
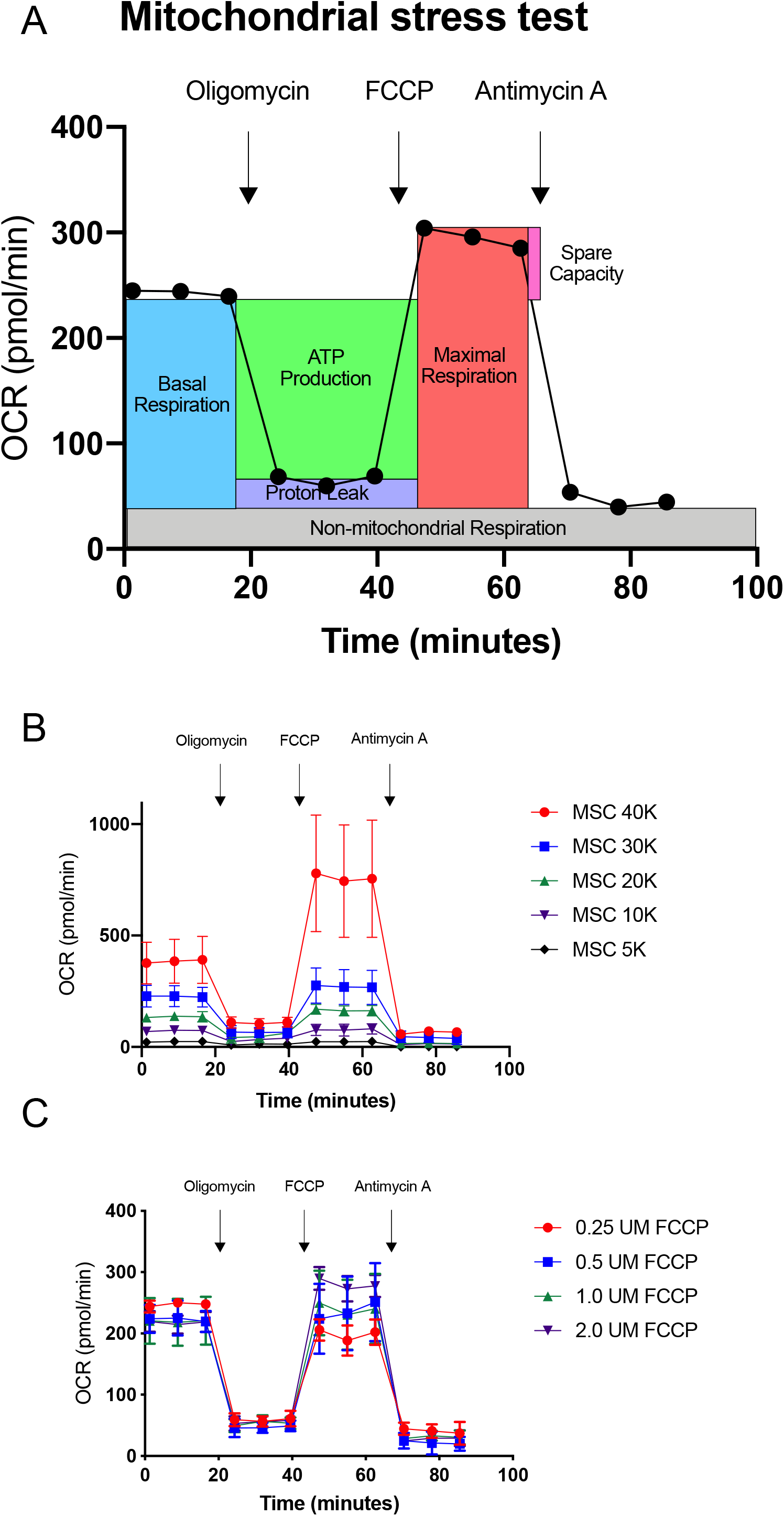
Optimization of the cell number and FCCP concentration for the mitochondrial stress test using Seahorse technology. (A) The oxygen consumption rate (OCR) in MSCs was measured using Seahorse technology followed by subsequent measurements after injection of mitochondrial toxins: oligomycin, FCCP, and antimycin A. Using the mitochondrial stress test Basal OCR, ATP production, Maximum OCR, Spare capacity, Nonmitochondrial respiration and Proton leak were determined. (B) Mitochondrial stress test with different MSCs densities per well (5K, 10K, 20K, 30K and 40K) using 1.0 μM FCCP. (C) Mitochondrial stress test with 30K MSCs per well using different concentrations FCCP (0.25, 0.5, 1.0 and 2.0 μM). N=5-7, 1 donor with 5-7 replicates per donor. Graphs represent mean with SD.

**Fig EV5.Related to Fig3.**
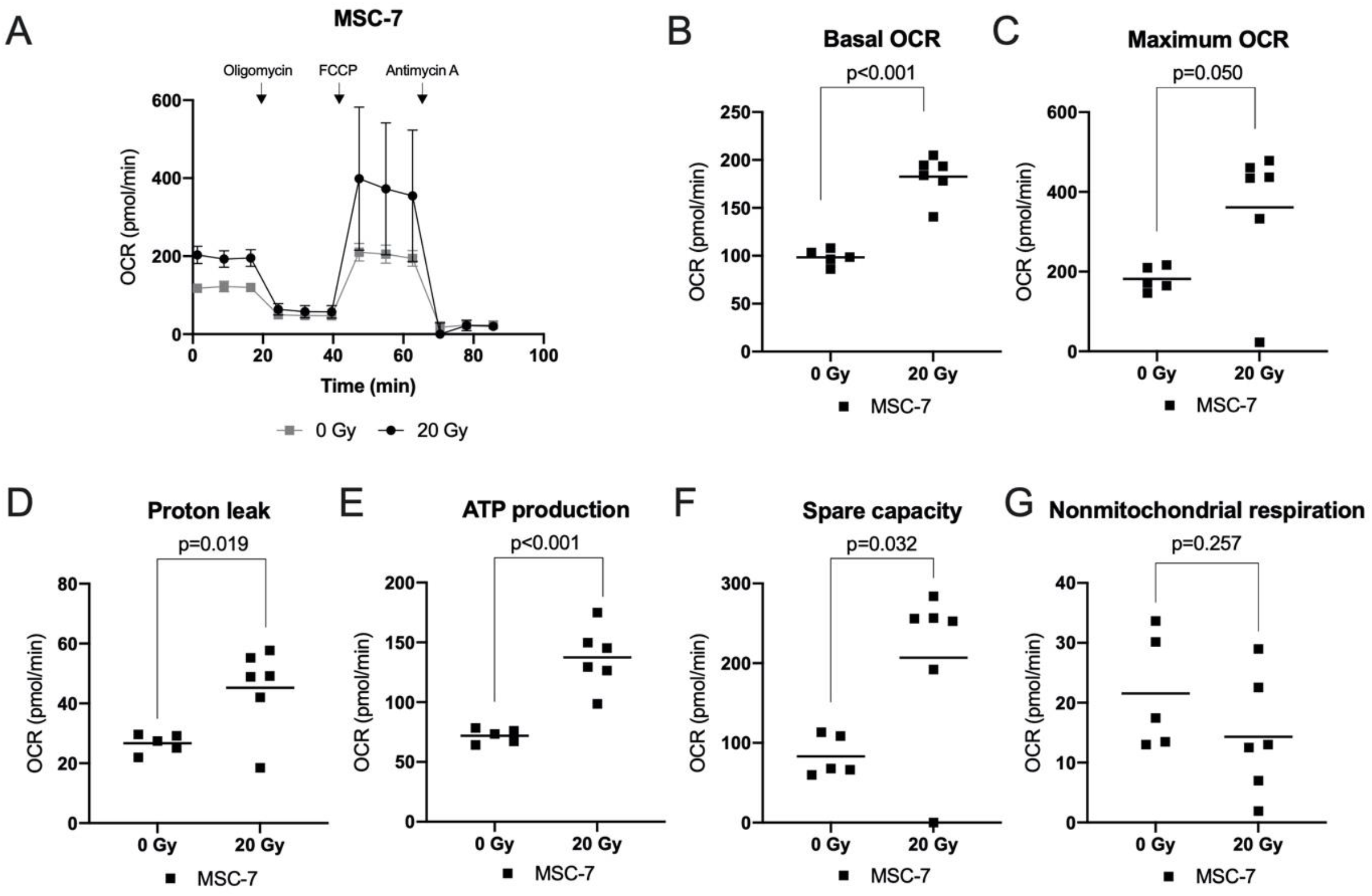
Increased oxygen consumption rate (OCR) in TWIST1 silenced MSCs. (A) Graph shows the OCR in MSCs irradiated with 0 or 20 Gy after addition of oligomycin, FCCP and antimycin A. Values represent mean with SD, N=5-6 replicates. (B-G) Graphs show calculated values for basal OCR (B), maximum OCR (C), proton leak (D), ATP production (E), spare capacity (F) and non-mitochondrial respiration (G) in MSCs irradiated with 0 or 20 Gy. N=5-6 replicates, unpaired t-test. Graphs show individual data points and mean.

**Fig EV6.Related to Fig4.**
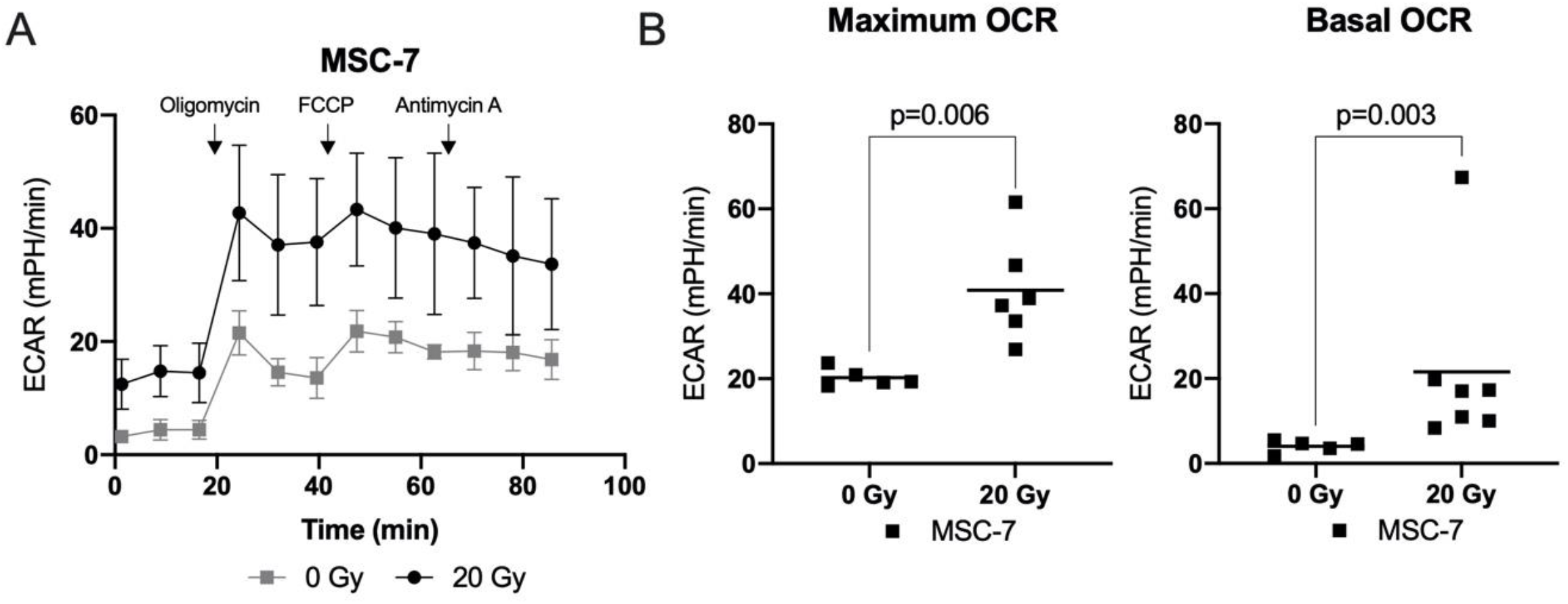
Increased extrcellular acidification rate (ECAR) in irradiated MSCs. (A) Graph shows the ECAR in MSCs irradiated with 0 or 20 Gy after addition of oligomycin, FCCP and antimycin A. Values represent mean with SD, N=5-6 replicates. (B) Graphs show ECAR values for basal OCR and maximum OCR in MSCs irradiated with 0 or 20 Gy. N=5-6 replicates, unpaired t-test. Graphs show individual data points and mean

**Table EV1.**
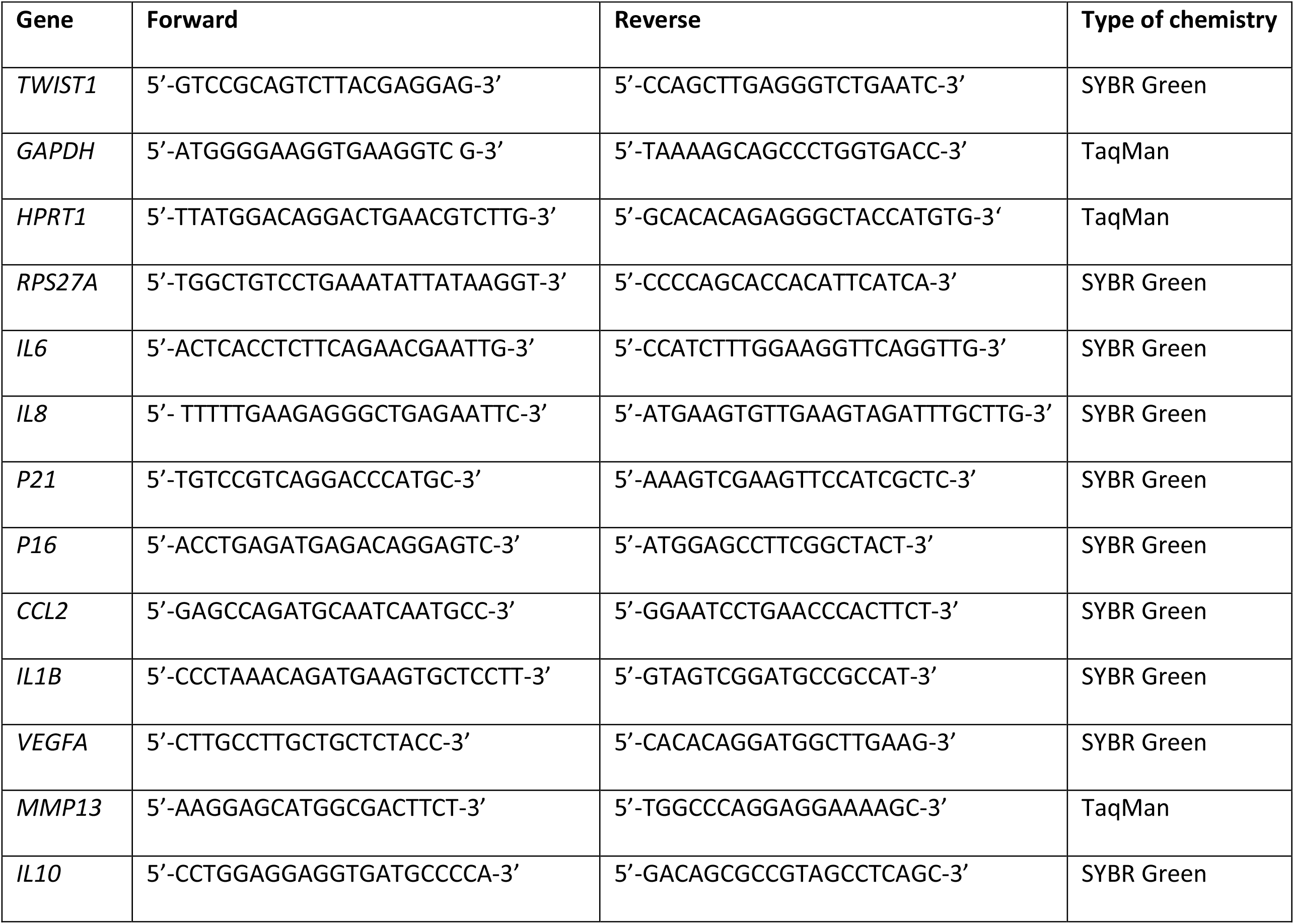
Primer sequences.

